# Clearance of DNA damage-arrested RNAPII is selectively impaired in Cockayne syndrome cells

**DOI:** 10.1101/2024.05.17.594644

**Authors:** Paula J. van der Meer, George Yakoub, Yuka Nakazawa, Tomoo Ogi, Martijn S. Luijsterburg

**Affiliations:** Department of Human Genetics, Leiden University Medical Center, Leiden, The Netherlands; Department of Genetics, Research Institute of Environmental Medicine, Nagoya University, Nagoya, Japan; Department of Human Genetics and Molecular Biology, Nagoya University Graduate School of Medicine, Nagoya, Japan

**Author notes:** Equal contribution (Paula J. van der Meer, George Yakoub). **Corresponding author:** Martijn S. Luijsterburg.

## Abstract

Arrest of elongating RNA polymerase II (RNAPII) at DNA lesions initiates transcription-coupled repair (TCR), involving the concerted action of specific TCR factors, followed by downstream nucleotide excision repair steps. Remarkedly, only congenital defects in the CSA or CSB genes cause the neurodegenerative disorder Cockayne syndrome, which is not observed with other TCR genes, despite their equal importance in TCR. An explanation for this discrepancy has been lacking. In this study, we developed an assay to track the fate of elongating RNAPII at sites of UV-induced DNA lesions. Employing this method on an isogenic collection of TCR knockout cells reveals a selective RNAPII clearance defect in cells defective in CSA or CSB, in contrast to knockouts of other TCR genes. Our findings provide evidence that a deficiency in RNAPII processing and prolonged transcription arrests in response to DNA damage, rather than compromised DNA repair, may underlie the Cockayne syndrome-like neurodegenerative phenotype.

## Introduction

During gene transcription, RNA polymerase II (RNAPII) translocates along the DNA, synthesizing a nascent transcript. DNA lesions in the template strand, including UV-induced cyclobutene–pyrimidine dimers (CPDs) and 6–4 pyrimidine–photoproducts (6-4PPs), cause stalling of elongating RNAPII and trigger a genome-wide transcriptional arrest^1,2^. Stalled RNAPII acts as the damage sensor to initiate transcription-coupled repair (TCR), a sub-pathway of nucleotide excision repair (NER) that specifically removes transcription-blocking lesions from actively transcribed DNA strands^3,4^.

DNA damage-arrested RNAPII is recognized by the Cockayne syndrome B (CSB) protein, a DNA-dependent ATPase that binds DNA upstream of RNAPII. Subsequently, CSB recruits CSA to stalled RNAPII, which is the substrate-recognition subunit of a DDB1-CUL4A-RBX1 ubiquitin ligase complex (CRL4^CSA^). Next, UV-stimulated scaffold protein A (UVSSA) is recruited to the complex in a CSA-dependent manner^5–7^. The RNAPII-bound transcription elongation factor ELOF1 plays an essential role in TCR complex assembly^8,9^. Following its recruitment, CSA docks onto RNAPII-bound ELOF1, with UVSSA further stabilizing the ELOF1-CSA interface^10^. Together, these events lead to the ubiquitylation of damage-stalled RNAPII at a single lysine in its largest subunit (RPB1-K1268)^8–12^.

The assembly of TCR factors together with RNAPII ubiquitylation culminates in the recruitment of transcription factor IIH (TFIIH) to RNAPII. TFIIH is a general repair factor that is shared between TCR and the global genome repair (GGR) sub-pathway of NER, in which damage-sensing protein XPC recruits TFIIH to the damage site^13,14^. Following recruitment, TFIIH is bound by XPA and endonucleases XPG and ERCC1-XPF, leading to the unwinding of the DNA and a dual incision around the damaged nucleotide^15–19^. These incisions release a 22– 30 nucleotide DNA fragment containing the lesion, which is followed by repair synthesis and ligation of the single-stranded DNA gap^19–21^.

Congenital defects in TCR genes can lead to widely varying clinical manifestations. Cockayne Syndrome (CS) patients, who carry bi-allelic mutations in CSA or CSB genes, suffer from severe progressive neurodegeneration and developmental abnormalities^22^. Conversely, individuals with the majority of UV-sensitive Syndrome (UV^S^S), caused by defective UVSSA, only display mild photosensitivity^6,23–26^. Defective XP proteins A-G cause Xeroderma Pigmentosum (XP), characterized by photosensitivity and cancer predisposition^27^. Remarkably, UV^S^S and most of XP do not cause the severe neurodegenerative phenotype of CS. However, it must be noted that rare cases with specific mutations in XPG or the TFIIH subunits XPB or XPD can lead to a combined XP/CS phenotype^28,29^. Additionally, distinct mutations in TFIIH subunits XPB, XPD, or TTDA can cause Trichothiodystrophy (TTD), which shares features with CS but also includes brittle hair, nails, and scaly skin^27^. Lastly, very rare mutations in genes encoding the multifunctional ERCC1-XPF can cause a complex combination of phenotypes depending on which functions are affected, including liver and kidney dysfunction, XP or Fanconi Anaemia, but also CS-like features^30,31^. Nevertheless, it appears that only congenital defects in the *CSA* or *CSB* genes result in the neurodegenerative disorder Cockayne syndrome, a phenomenon generally not observed with other TCR genes despite their equal importance and contribution to TCR.

An explanation for this discrepancy has been lacking. It has been suggested that it is not the TCR defect but the inability to process damage-stalled RNAPII that causes this neurodegenerative phenotype^6,32^. However, the precise regulation of RNAPII processing and the exact fate of damage-stalled RNAPII has remained somewhat ambiguous because the experimental tools to address this directly have been lacking^1,25,33,34^. Evidently, the stepwise recruitment of the early TCR proteins to damage-stalled RNAPII leads to the displacement and degradation of RNAPII that shields the DNA lesion, which is necessary to allow the downstream NER proteins access to the lesion^5^. During this process, RNAPII itself undergoes three main events upon stalling: i) ubiquitylation of RPB1 at K1268, ii) removal of RNAPII from the DNA lesion, iii) degradation of ubiquitylated RNAPII by the proteasome.

RNAPII processing, removal, and degradation have never been assessed systematically. Previous studies performed in primary fibroblasts derived from CS patients showed that ubiquitylation and degradation of RNAPII after UV damage were impaired in cells deficient in either CSA or CSB^6,32^. In contrast, primary cells deficient in UVSSA had accelerated RNAPII degradation after UV despite a defect in RNAPII ubiquitylation^6^. Somewhat confusingly, one study using immortalized human cells reached opposing conclusions. In that study, CSB knockout cells engineered to ectopically express a tagged version of RNAPII’s largest RPB1 subunit exhibited rapid and complete degradation of RNAPII after UV, leading to the conclusion that CS is associated with unstable RNAPII^35^. Because these studies were performed in cells with different genetic backgrounds, it is challenging to systematically evaluate the different phenotypes. Moreover, existing experimental tools that measure RNAPII stalling, such as TCR-seq^8^, do not distinguish between prolonged stalling of RNAPII on a lesion and recurrent RNAPII arresting on unrepaired lesions.

Here, we developed an approach to specifically measure the clearance of DNA damage-arrested RNAPII. In combination with the ubiquitylation and degradation of RNAPII upon UV, we investigated a collection of seven isogenic knockout cell lines of key TCR factors to uncover which of these TCR factors drive the processing and determine the fate of damage-stalled RNAPII.

## Results

### Local DRB run-off: a method to track DNA damage-stalled RNAPII clearance in situ

The fate of RNAPII stalled at DNA lesions is unclear, largely due to the unavailability of methods to directly visualize this process. Therefore, we developed an assay to first induce, on average, one local UV damage per nucleus using microfilters, then track the fate of RNAPII and measure its potential clearance at these sites of local UV damage. To specifically measure the local levels of already elongating RNAPII without the interference of multiple iterations of new elongating RNAPII arresting on unrepaired lesions, we added to the cells the transcription elongation inhibitor 5,6-dichloro-1-β-D-ribofuranosylbenzimidazole (DRB) directly after UV treatment. DRB inhibits the transition of new RNAPII complexes from the initiation (RNAPIIa) to the hyperphosphorylated elongation (RNAPIIo/S2) form. Finally, the clearance of damage-stalled RNAPII was monitored at different time points through immunofluorescence (IF) staining of elongating RNAPII-S2 (Fig 1a; Fig S1). By performing this assay in RPE1-hTERT WT cells, we detected a gradual decrease in RNAPII-S2 following DRB treatment outside of the locally damaged area in the nucleus within a time course of 2 h (Fig 1b). This reduction is consistent with elongating RNAPII (travelling ∼120 kb in 1 h), eventually running off genes and becoming de-phosphorylated. However, at sites of local UV damage, marked by CPD staining, we observed an accelerated loss (∼2-fold) of RNAPII-S2 compared to the signal outside of the damaged area, reflecting the active clearance of DNA damage-stalled RNAPII from chromatin (Fig 1b-c; Fig S1b-c). Altogether, we developed a new assay capable of directly assessing the clearance of stalled RNAPII from DNA lesions.

**Fig 1.**
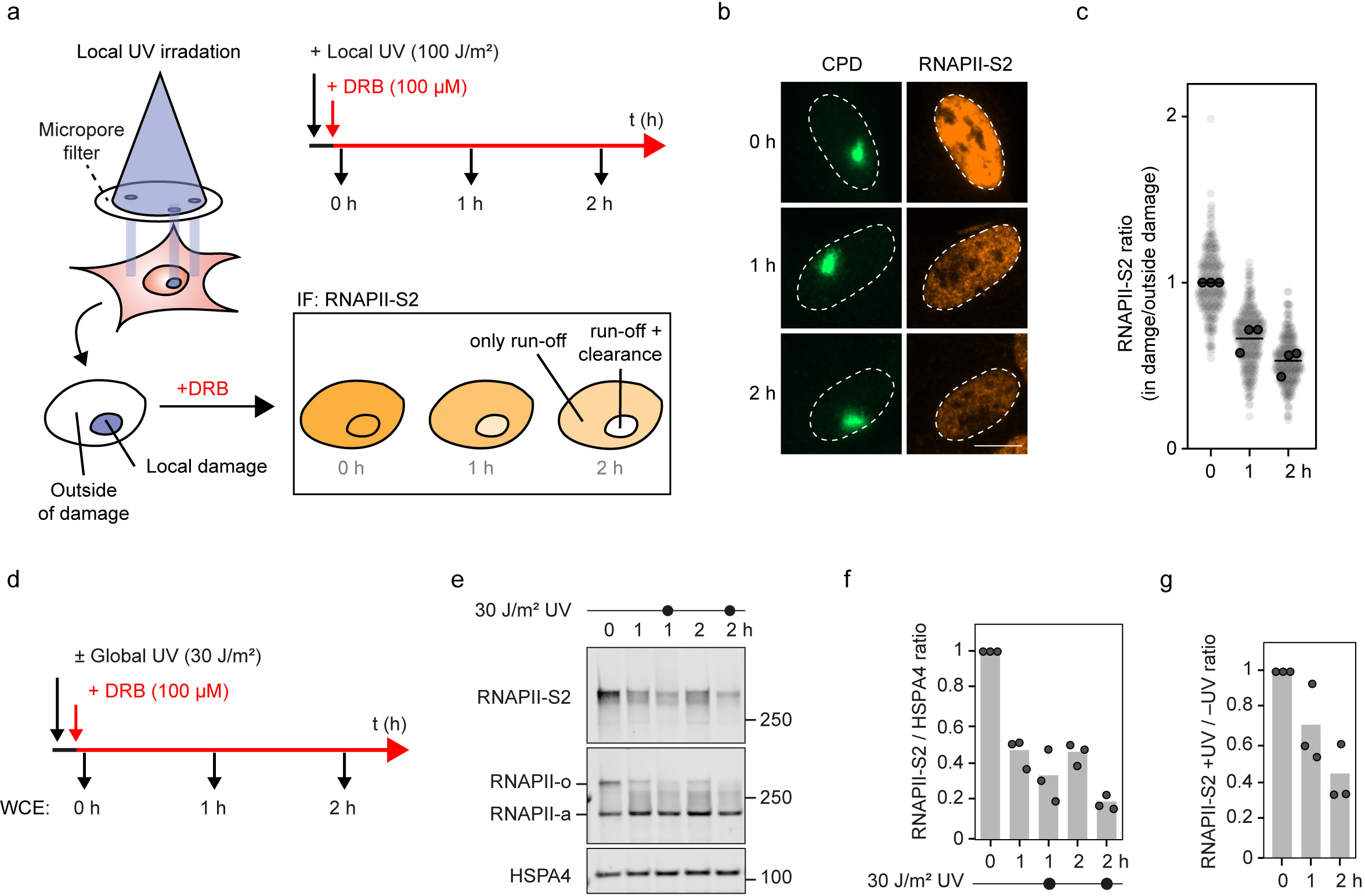
Experimental approach to measure local and global DRB run-off of elongating RNAPII. **a** Outline of the local DRB run-off assay by immunofluorescence imaging. **b.** Representative images of RNAPII-S2 signal in RPE1 WT cells at 0, 1, and 2 h after local UV damage (100 J/m^2^) marked by CPD. Cells were treated with 100 μM DRB immediately after UV irradiation to prevent new RNAPII molecules from moving into gene bodies: scale bar, 10 μm. **c.** Quantification of the images in **b**. The RNAPII-S2 mean pixel intensity inside the local damage was divided by the mean pixel intensity outside the local damage in the same cell nucleus to correct for a decrease of RNAPII-S2 signal due to a general RNAPII run-off caused by DRB. A ratio below 1 signifies that RNAPII-S2 loss in the local damage is faster than the general run-off outside the local damage. All cells are depicted as individual semi-transparent data points. The bar represents the mean of all data points. The individual means of three biological replicates are depicted as solid circles with black strokes. **d.** Outline of the global DRB run-off assay by western blotting. **e.** A representative image of the RNAPII signal detected by western blot from whole cell lysates of RPE1 WT cells at 0, 1, and 2 h after global UV damage (30 J/m^2^). Cells were treated with 100 μM DRB immediately after UV irradiation to prevent new RNAPII molecules from moving into gene bodies. An antibody against phosphorylated RNAPII detects only the elongating form RNAPII-S2 (upper blot). An antibody targeted against the N-terminal domain of RNAPII detects both initiating (RNAPIIa) and the elongating (RNAPIIo) forms (middle blot). Note that RNAPII-S2 and RNAPIIo represent the same pool of phosphorylated RNAPII. Different abbreviations were used to distinguish between the signals detected by the different antibodies. The HSPA4 signal was used as a loading control (lower blot). **f.** Quantification of the blots in **e**. The RNAPII-S2 mean pixel intensity was divided by the mean pixel intensity of HSPA4. Then, the value of each time point and condition was normalized to 0 h. A ratio below 1 in unirradiated cells indicates a general RNAPII-S2 run-off caused by DRB. The additional drop in the ratio in UV-treated cells indicates an active RNAPII-S2 degradation by TCR. The bar represents the mean of three independent biological replicates. The individual means of the three biological replicates are depicted as solid circles with black strokes. **g.** Quantification of the blots in **e**. The RNAPII-S2 mean pixel intensity was divided by the mean pixel intensity of HSPA4. Then, the value of each UV-treated time point was normalized to its respective unirradiated condition. Here, a ratio below 1 indicates an active RNAPII-S2 degradation by TCR. The bar represents the mean of three independent biological replicates. The individual means of the three biological replicates are depicted as solid circles with black strokes.

### Global DRB run-off: a method to detect DNA damage-stalled RNAPII degradation

Although the local DRB run-off assay can measure the clearance of elongating RNAPII from UV lesions, it runs short in determining the fate of RNAPII following its eviction. Therefore, we adjusted our DBR run-off approach to investigate RNAPII degradation. To this end, cells were globally exposed to UV and incubated in DRB-containing media. After 1 and 2 h intervals, the levels of RNAPIIo/S2 were analyzed from whole cell extracts by SDS-PAGE and western blotting (Fig 1d). Similar to the local IF approach, DRB treatment alone without any UV damage caused a gradual reduction in RNAPIIo/S2 levels, indicating that elongating RNAPII ran off gene bodies and became de-phosphorylated (Fig 1e-f). Interestingly, UV-treated samples after DRB showed a faster reduction in RNAPIIo/S2 relative to unirradiated samples treated with DRB, indicative of active degradation upon eviction from DNA lesions (Fig 1e-g). Taken together, we established two parallel approaches to specifically monitor the clearance and degradation of elongating RNAPII upon encountering UV-induced DNA damage.

### Generating and validating an isogenic collection of TCR-defective cell lines

To investigate the involvement of individual TCR factors in the active clearance and degradation of lesion-stalled RNAPII, we generated a collection of seven isogenic TCR-gene knockouts in human RPE1-hTERT cells. The collection included cells with genetic inactivation of early TCR factors (CSB, CSA), factors that stabilize the RNAPII-bound TCR complex and recruit TFIIH (ELOF1, UVSSA), and downstream NER factors that act after RNAPII displacement to repair the actual lesions (XPA, XPG, ERCC1) (Fig 2a). The efficient deletion of most TCR factors was confirmed by western blot analysis, showing the loss of the respective TCR proteins (Fig 2b, S2a-b). Knockouts of ELOF1 and UVSSA were validated by Sanger sequencing because commercially available antibodies do not recognize these endogenous proteins in RPE1 cells (Fig 2c). This unique isogenic knockout (KO) collection enables a side-by-side comparison of how different factors at various stages of TCR affect RNAPII clearance and degradation.

**Fig 2.**
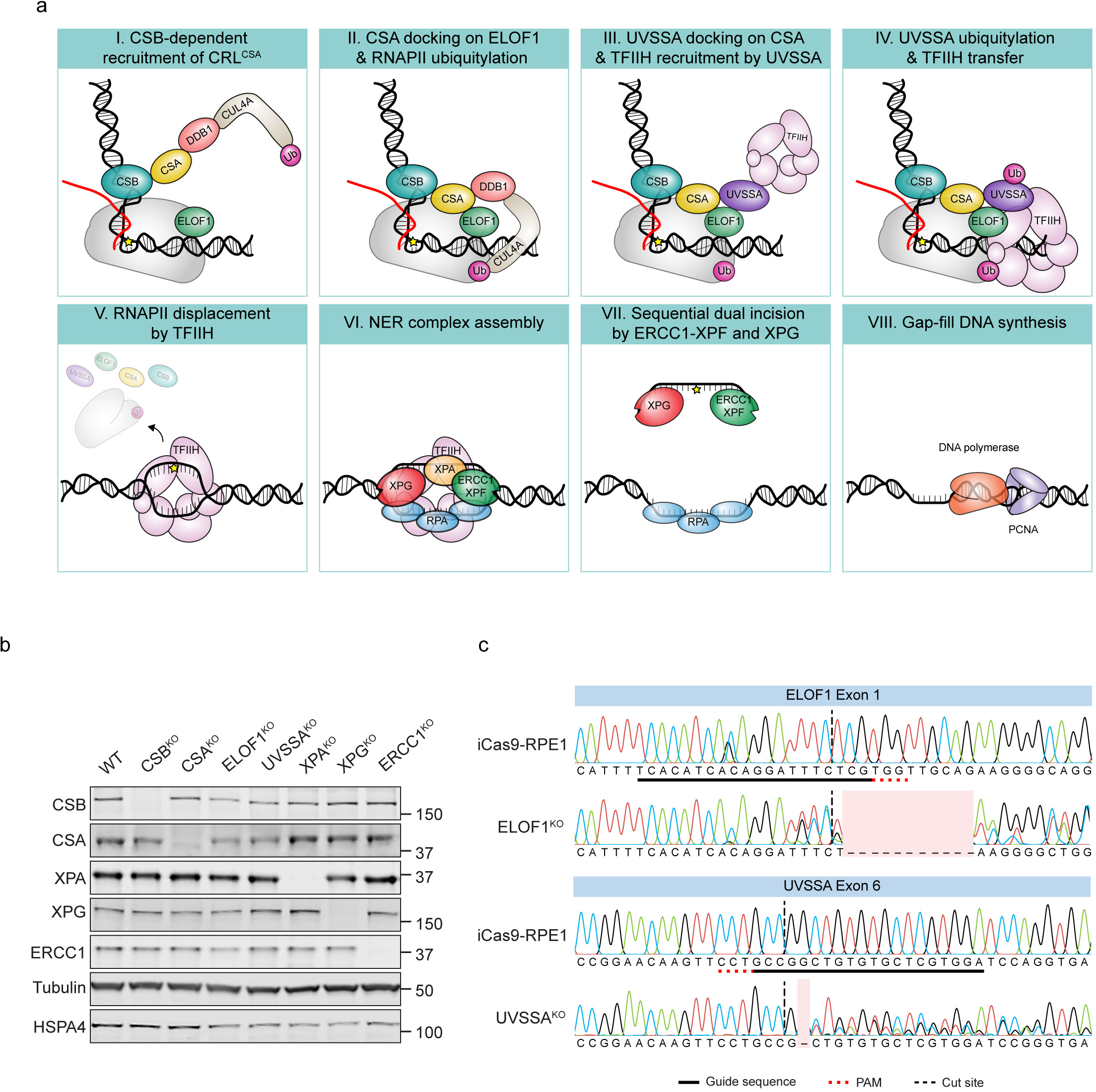
Generation and validation of isogenic collection of TCR knockout cells. **a.** A cartoon depicting the molecular mechanism of TCR and RNAPII processing. Each step is labeled with a Latin Numeric. **b.** Validation of the TCR knockouts by western blotting. The indicated antibodies were used to analyze whole cell lysates from parental and knockout RPE1 cell lines. **c.** Validation of ELOF1^KO^ and UVSSA^KO^ cell lines by Sanger sequencing. Chromatographs of the genomic regions of the respective genes expanding upstream and downstream of the cut site by the sgRNA.

### TCR-dependent incisions are reduced in TCR-defective cell lines

Among the final steps in TCR is the dual incision that excises the damaged DNA strand, executed by the endonucleases XPG and ERCC1-XPF^1^. The chemotherapeutic trabectedin is specifically toxic to cells with proficient TCR because it generates RNAPII-stalling lesions that are processed by TCR proteins but block the 3’ incision by XPG. The resulting uncoupled 5’ incision by ERCC1-XPF creates a persistent single-strand break that is converted into a double-strand break (DSB) during replication^36,37^ (Fig 3a). However, these single-strand and following DSBs are not generated when TCR is defective^10^. To demonstrate that TCR is impaired in each of the KO cell lines in this study, we employed a repair assay that we recently established^10^ and tested to what extent these TCR-mediated incisions and DSBs were generated. We treated cells with trabectedin and measured γH2AX levels in dividing (EdU-positive) cells after four hours. Indeed, WT cells show elevated γH2AX levels upon trabectedin treatment, indicating that TCR is proficient in these cells (Fig 3b-c). However, all seven KO cell lines had reduced γH2AX levels, in agreement with their expected TCR defect (Fig 3b-c). Taken together, we demonstrate that all KO cell lines used in this study are defective in TCR.

**Fig 3.**
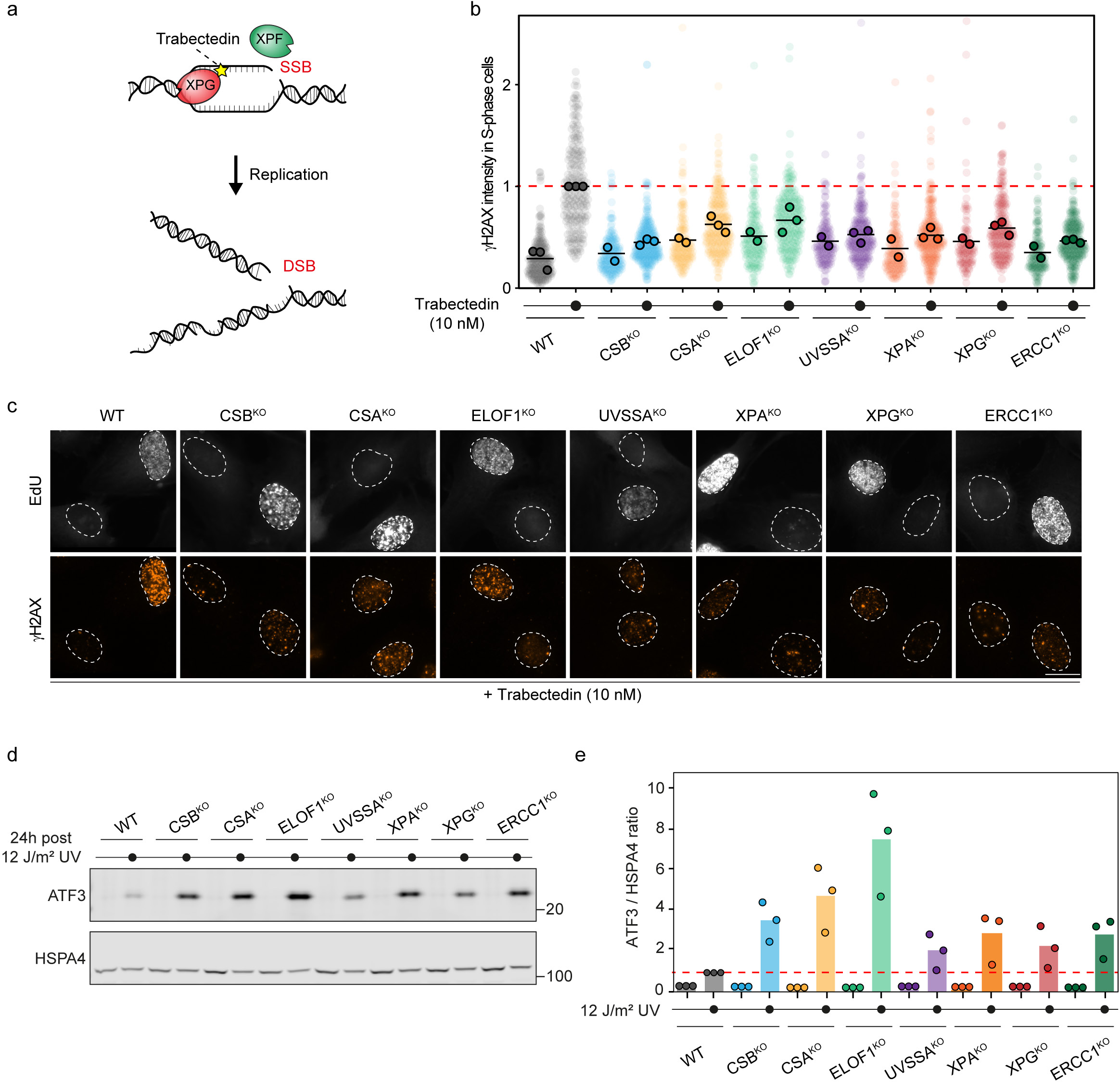
The isogenic knockout collection is defective in TCR and transcription recovery. **a.** Model of TCR-dependent double-strand break formation in replicating cells resulting from trabectedin treatment. **b.** Quantification of TCR-dependent γH2AX induction following trabectedin exposure in replicating cells labeled with 5-ethynyl-deoxyuridine (EdU) in RPE1 cells with the indicated genotype. The γH2AX levels were normalized to the average of the trabectedin-treated parental WT cells within each experiment. All cells are depicted as individual semi-transparent data points. The bar represents the mean of all data points. The individual means of three biological replicates are depicted as solid circles with black strokes. **c.** Representative images of **b**: scale bar, 10 μm. **d.** A representative image of ATF3 protein levels detected by western blot from whole cell lysates of RPE1 WT and the indicated isogenic TCR knockouts. Cells were either kept unirradiated or exposed to 12 J/m^2^ UV-C and collected after 24 h. The HSPA4 signal was used as a loading control. **e.** Quantification of the blots in **d**. The ATF3 mean pixel intensity was divided by the mean pixel intensity of HSPA4 to correct for loading. Then, the value of each time point and condition was normalized to the value of UV-treated parental WT cells. The bar represents the mean of three independent biological replicates. The individual means of the three biological replicates are depicted as solid circles with black strokes.

### Recovery from transcription inhibition is impaired in TCR-defective cell lines

The presence of UV-induced DNA damage can induce a global transcription shutdown that gradually recovers within 24 h in WT cells. However, CSB- or CSA-deficient cells fail to recover transcription of approximately ∼5000 genes due to defective TCR, culminating in the constitutive presence of the transcription repressor ATF3 at gene promoters^38^. Hence, a prolonged high level of ATF3 is considered another hallmark of defective TCR. To investigate whether each of the TCR knockout cell lines is impaired in transcription recovery, we measured the protein levels of ATF3 in UV-exposed and undamaged conditions using western blotting. In WT cells, ATF3 levels were slightly upregulated in response to UV damage. However, all TCR-deficient cell lines showed at least a 2-fold increase in ATF3 levels 24 h after UV irradiation (Fig 3d-e, S2c-d). Notably, ATF3 upregulation in CSB^KO^, CSA^KO^, and especially ELOF1^KO^ was far more pronounced than in the other knockout cell lines. This observation correlates well with the previously demonstrated transcription recovery profile of these cells, where WT cells recover from transcription inhibition within 24 h, indicated by the low ATF3 levels, whereas transcription does not restart properly in TCR-deficient cells, as illustrated by the prolonged high ATF3 levels in these cells after UV treatment. While we have previously observed increased ATF3 levels in UVSSA^KO^ cells^8^, the inability of XPA^KO^, XPG^KO^, and ERCC1^KO^ cells to downregulate ATF3 and restore transcription was not reported before. Altogether, these results indicate that similar to CS cells, the knockout cell lines of other key TCR factors are also defective in recovering from transcription arrest.

### The impact of key TCR factors on the clearance of DNA damage-stalled RNAPII

Having confirmed that each of the knockout cell lines in our collection is defective in TCR, we used the local DRB run-off approach to track RNAPII clearance in the TCR knockout cell lines (Fig 4a). Due to the DRB treatment, the levels of elongating RNAPII outside of the locally damaged area in the nucleus decreased gradually within a time course of 2 h, as was observed before. While WT cells displayed an accelerated loss of RNAPII-S2 signal at the sites of local UV damage, the ratio between RNAPII-S2 signal inside and outside of the local damage in CSB^KO^ and CSA^KO^ cells stayed high, clearly reflecting a defect in the removal of stalled RNAPII-S2 from DNA lesions (Fig 4b-c; also see Fig S1c-d). In ELOF1^KO^ and UVSSA^KO^ cells, removal of damage-stalled RNAPII was attenuated, exhibiting a phenotype between that observed in WT and CS cells. Interestingly, the clearance of damage-arrested RNAPII in XPA^KO^, XPG^KO^, and ERCC1^KO^ cells was similar to that in WT cells. Hence, based on our new assay, CS and non-CS cells differ in their ability to clear RNAPII-S2 from UV damage sites.

**Fig 4.**
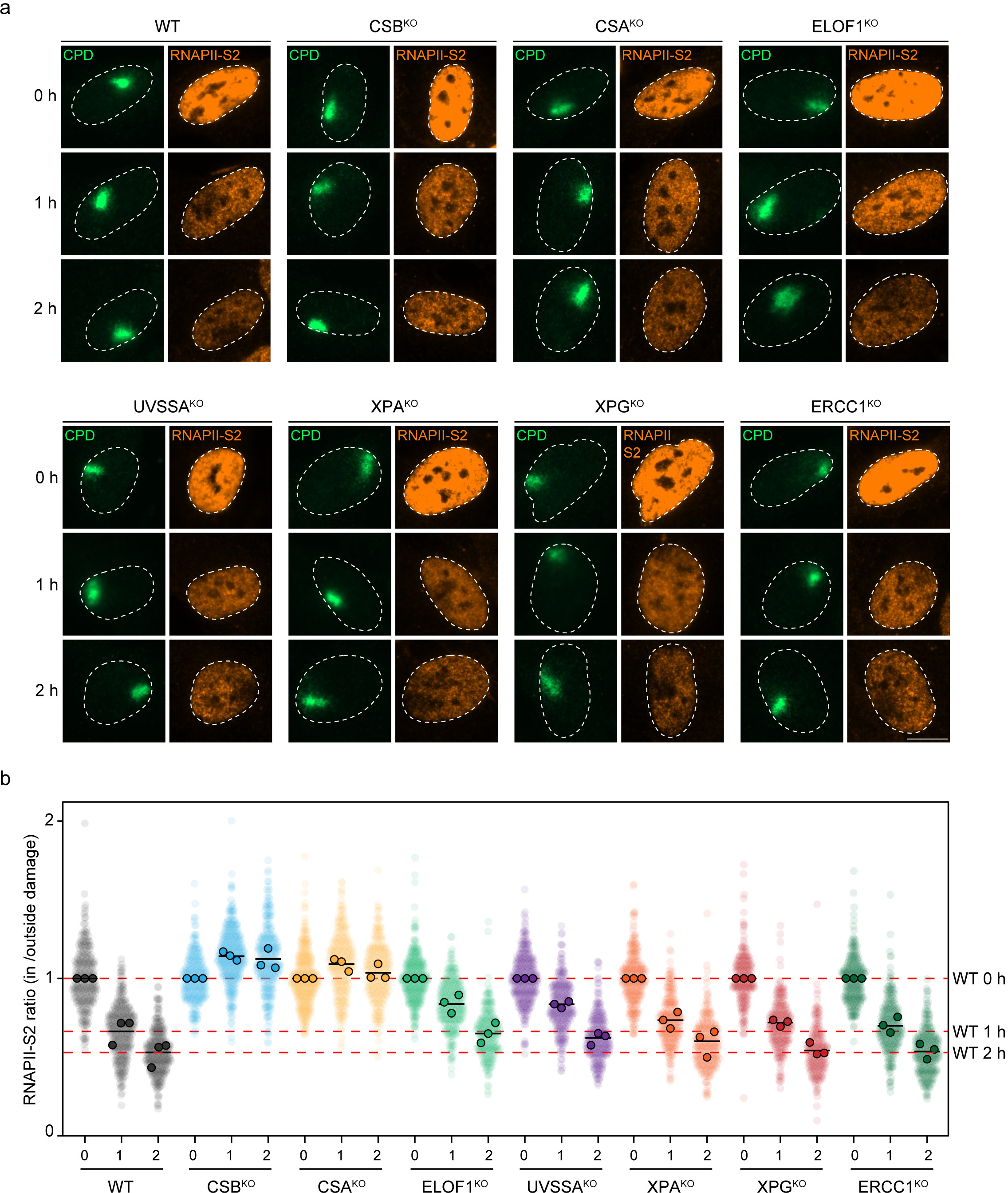
Local clearance of damage-stalled RNAPII in TCR knockout cells. **a.** Representative images of RNAPII-S2 signal in RPE1 cells with the indicated genotype at 0, 1, and 2 h after local UV damage (100 J/m^2^) marked by CPD. Cells were treated with 100 μM DRB immediately after UV irradiation to prevent new RNAPII molecules from moving into gene bodies: scale bar, 10 μm. **b.** Quantification of the images in **a**. The RNAPII-S2 mean pixel intensity inside the local damage was divided by the mean pixel intensity outside the local damage in the same cell nucleus to correct for a decrease of RNAPII-S2 signal due to a general RNAPII run-off caused by DRB. A ratio below 1 signifies that RNAPII-S2 loss in the local damage is faster than the general run-off outside the local damage. All cells are depicted as individual semi-transparent data points. The bar represents the mean of all data points. The individual means of three biological replicates are depicted as solid circles with black strokes.

### The impact of key TCR factors on degradation of DNA damage-stalled RNAPII

To confirm that the clearance of elongating RNAPII from UV lesions is manifested in its degradation, we repeated the global DRB run-off assay in the seven isogenic TCR knockout cell lines (Fig 1d). The cells were globally exposed to UV damage and incubated in DRB-containing media. Similar to the local IF approach, DRB treatment alone without any UV damage caused a gradual reduction in RNAPIIo/S2 levels in all tested TCR knockouts, suggesting that elongating RNAPII ran off gene bodies, while RNAPIIa was still clearly present at promoters due to the DRB treatment (Fig 5a-b, S3a-d). In contrast, UV-treated samples showed a faster reduction of RNAPIIo/S2 in WT cells, suggesting its active degradation due to TCR (Fig 5a-d, S3a,c, S4a-d). Interestingly, only CSB^KO^ and CSA^KO^ cells, but none of the other TCR knockouts, displayed significantly reduced degradation of RNAPII-S2 after UV damage (Fig 5a-d, S3a,c, S4a-d). This is especially clear when comparing the RNAPIIo signal between UV-irradiated and unirradiated cells treated with DRB (Fig 5a). Together with the local DRB run-off results, the RNAPII clearance and degradation observed in CSB^KO^ and CSA^KO^ cells correlate well with the severe CS phenotype.

**Fig 5.**
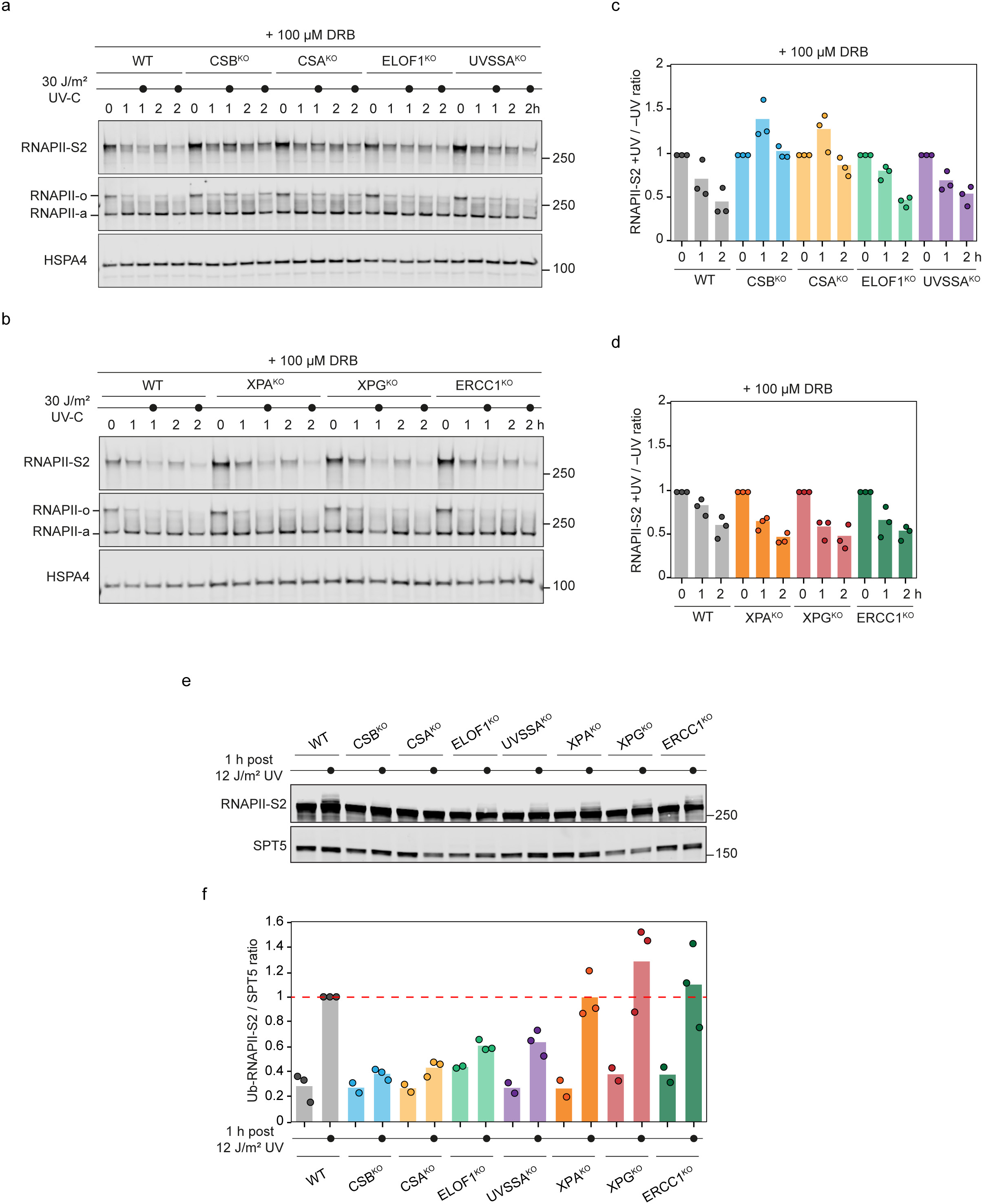
Ubiquitylation and global degradation of damage-stalled RNAPII in TCR knockout cells. **a-b.** Representative images of the RNAPII signal detected by western blot analysis from whole cell lysates of RPE1 WT and the indicated TCR knockout cells at 0, 1, and 2 h after global UV damage (30 J/m^2^). Cells were treated with 100 μM DRB immediately after UV irradiation to prevent new RNAPII molecules from moving into gene bodies. An antibody against phosphorylated RNAPII detects only the elongating form RNAPII-S2 (upper blot). An antibody targeted against the N-terminal domain of RNAPII detects both initiating (RNAPIIa) and the elongating (RNAPIIo) forms (middle blot). Note that RNAPII-S2 and RNAPIIo represent the same pool of phosphorylated RNAPII. Different abbreviations were used to distinguish between the signals detected by the different antibodies. The HSPA4 signal was used as a loading control (lower blot). **c-d.** Quantification of the blots in **a-b**. The RNAPII-S2 mean pixel intensity was divided by the mean pixel intensity of HSPA4. Then, the value of each UV-treated time point was normalized to its respective unirradiated condition. Here, a ratio below 1 indicates an active RNAPII-S2 degradation by TCR. The bar represents the mean of three independent biological replicates. The individual means of the three biological replicates are depicted as solid circles with black strokes. **e.** A representative image of RNAPII-S2 ubiquitylation detected by western blot analysis from whole cell lysates of RPE1 WT and the indicated TCR knockouts. Cells were treated with 12 J/m^2^ UV-C and collected after 1 h. The SPT5 signal was used as a loading control. **f.** Quantification of the blots in **e**. The RNAPII-S2 mean pixel intensity of the higher traveling bands, which are the ubiquitylated forms of RNAPII-S2, was divided by the mean pixel intensity of SPT5 to correct for loading. Then, the value of each time point and condition was normalized to the value of UV-treated parental WT cells. The bar represents the mean of three independent biological replicates. The individual means of the three biological replicates are depicted as solid circles with black strokes.

### UV-induced ubiquitylation of RNAPII in TCR-defective cell lines

When RNAPII arrests on a lesion, its largest subunit is ubiquitylated on a single lysine (RPB1-K1268) by CSB-mediated recruitment of the CRL4^CSA^ E3 ligase complex and subsequent docking on ELOF1 that is further stabilized by UVSSA to direct the ubiquitylation^10,11^. TCR-associated ubiquitylation is essential for efficient repair and for efficient RNAPII degradation following UV irradiation. To examine side-by-side to what extent the TCR-defective cell lines are proficient in UV-induced ubiquitylation of RNAPII, we irradiated cells and analyzed the ubiquitylation of chromatin-bound RNAPII. As expected, we observed a higher migrating band in WT cells upon UV irradiation, indicating efficient RNAPII ubiquitylation. In line with their known role in RNAPII ubiquitylation, CSB^KO^ and CSA^KO^ cells did not show any RPB1 ubiquitylation, reflecting their defects in RNAPIIo/S2 clearance and degradation at UV damage sites (Fig 5e-f). Interestingly, RPB1 ubiquitylation was strongly reduced but not completely abolished in ELOF1^KO^ and UVSSA^KO^, correlating with their slower RNAPII clearance rate. Knockouts of the factors that act downstream in the TCR pathway (XPA^KO^, XPG^KO^, and ERCC1^KO^) did not show any defect in RPB1 ubiquitylation (Fig 5e-f). Together, these data indicate that the level of RPB1 ubiquitylation correlates with the ability of cells to clear and degrade RNAPII-S2.

### TCR-defective patient cells manifest RNAPII ubiquitylation and degradation defects

To investigate if the knowledge gained from our new assays and isogenic cell lines might be reflected and beneficial in dissecting the disease phenotypes of TCR-deficient patients, we expanded our global UV damage assay to analyze RNAPII ubiquitylation and degradation in patient-derived cell lines. To this end, we treated seven primary fibroblast lines from patients with congenital TCR deficiency, along with a WT control, with cycloheximide to inhibit the synthesis of new RPB1, in order to better visualize RNAPII ubiquitylation and degradation profiles in primary fibroblasts^6^. Using antibodies recognizing elongating RNAPIIo/S2, we detected RNAPII ubiquitylation and degradation during a time-course of 6 h in WT fibroblasts (Fig 6a). Primary fibroblasts from individuals with CSB or CSA deficiency showed reduced RNAPII ubiquitylation and impaired processing and degradation of RPB1 (Fig 6a,c). While RNAPII ubiquitylation was attenuated in primary fibroblasts from a UVSSA patient, we still detected RNAPII processing and degradation, which appeared even faster than in WT cells, as we previously reported^6^. In contrast, primary fibroblasts obtained from patients with deficiencies in XPA, XPG, ERCC1, or XPF all displayed normal ubiquitylation and degradation of RNAPII, which again occurred even faster than in WT cells (Fig 6b-c, S5a-e). The results obtained using patient-derived primary fibroblasts match well with our findings using isogenic TCR knockout cells and reveal that the clearance of DNA damage-arrested RNAPII is selectively impaired in Cockayne syndrome cells.

**Fig 6.**
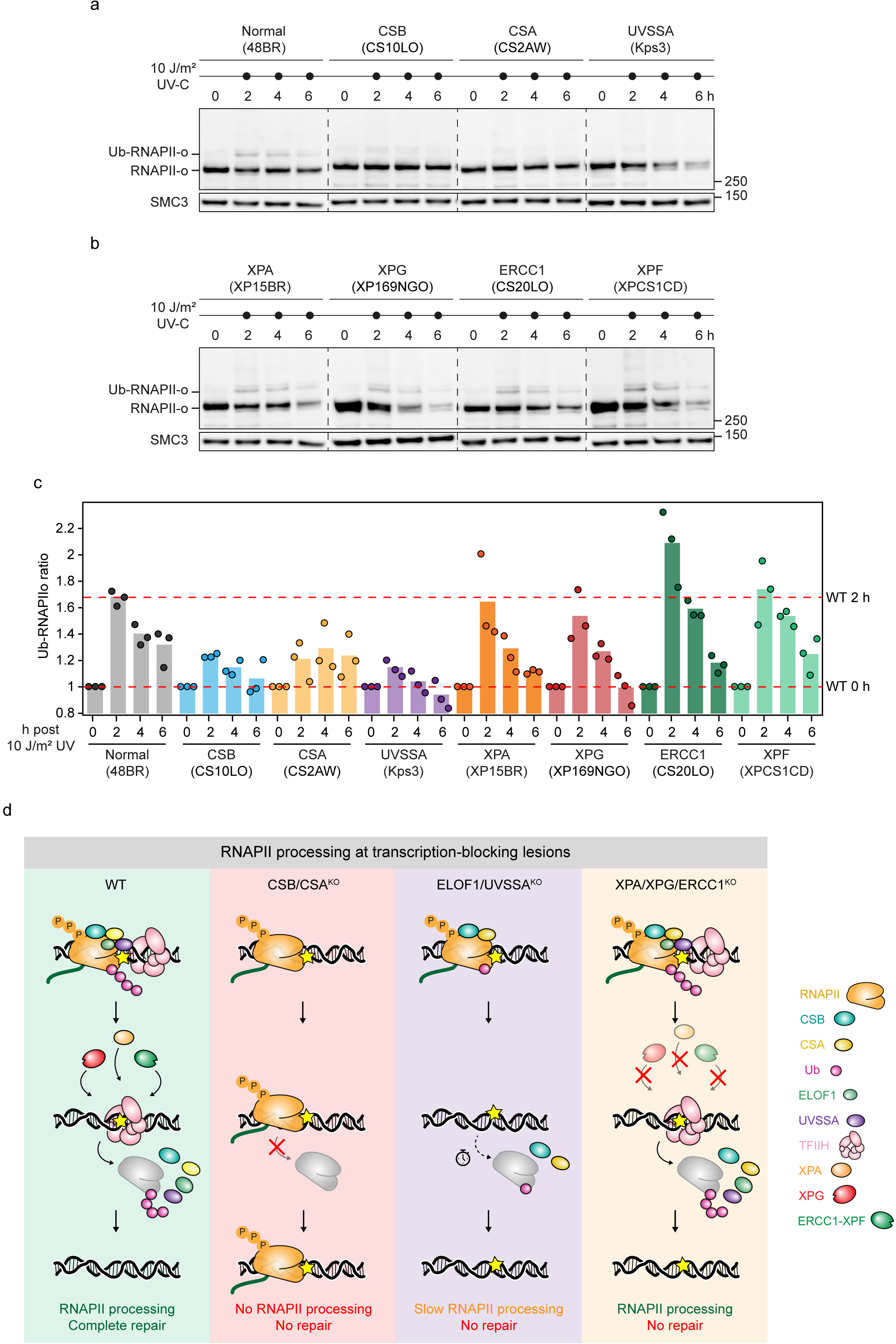
Ubiquitylation and global degradation of damage-stalled RNAPII in patient-derived cells. **a-b.** Representative images of the RNAPII signal detected by western blot analysis from whole cell lysates of patient-derives normal and the indicated TCR mutant cells at 0, 2, 4, and 6 h after global UV damage (10 J/m^2^). Cells were treated with 100 μM cycloheximide 1h before UV irradiation and incubated in the same media after UV exposure. An antibody against hyperphosphorylated C-terminal domain (CTD) Ser2/5 (H5) was used to detect the elongating RNAPIIo. The SMC3 signal was used as a loading control. **c.** Quantification of the blots in **a-b**. The Ub-RNAPIIo mean pixel intensity was divided by the mean pixel intensity of SMC3. Then, the value of each UV-treated time point was normalized to its respective 0 h condition. Here, a ratio below 1 indicates an active RNAPII-S2 degradation by TCR. The bar represents the mean of three independent biological replicates. The individual means of the three biological replicates are depicted as solid circles with black strokes. **d.** Graphical abstract of RNAPII processing in WT, non-CS, and CS cells. Defects in RNAPII processing and prolonged transcription arrest after DNA damage, rather than compromised DNA repair, may underlie the CS neurodegenerative phenotype.

## Discussion

Stalling of elongating RNAPII at DNA lesions initiates the orchestrated and sequential assembly of specific TCR factors, followed by downstream nucleotide excision repair to remove the lesion. Bi-allelic inactivating mutations in the *CSA* or *CSB* genes, the initial TCR factors that detect stalled RNAPII, cause the neurodegenerative disorder Cockayne syndrome. Strikingly, these devastating neurodegenerative symptoms are not observed when other essential TCR genes are inactivated despite their equal importance in repair. An explanation for this discrepancy was primarily lacking due to three major shortcomings: i) the absence of experimental tools, ii) divergent explanations for the underlying causes of the CS phenotype, and iii) the broad heterogeneity in the genetic backgrounds of the studied patient samples and cellular models.

In this study, we generated an isogenic collection of TCR knockout cell lines and developed an experimental setup to track the fate of elongating RNAPII upon stalling at UV-induced lesions. This new experimental tool enabled us to measure the clearance of damage-stalled RNAPII in a time-resolved manner. In parallel, we assessed the UV-induced ubiquitylation and degradation of RNAPII in the isogenic TCR knockouts. Combining these different approaches gave us the unique opportunity, for the first time, to directly compare side-by-side the impact of key TCR factors on the three aspects of RNAPII processing. Interestingly, we discovered that only CSB and CSA knockout cells have defects in UV-induced ubiquitylation of stalled RNAPII, its clearance from damage sites, and degradation (Fig 6d). In contrast, the rest of the isogenic TCR knockouts showed mild (ELOF1^KO^ and UVSSA^KO^) or no defect (XPA^KO^, XPG^KO^, and ERCC1^KO^).

The new RNAPII clearance assay presented in this study is widely applicable in many laboratory settings. Besides the side-by-side comparison of isogenic knockout cell lines presented in this study, this assay could be used, for example, in a clinical setting to assess RNAPII clearance in patient cell lines for the diagnosis of CS. Moreover, this method may be used to further investigate TCR or characterize RNAPII clearance in response to other types of DNA damage, provided that creating local DNA damage is possible. We envision an application in studying transcription responses to DNA double-strand breaks, for which ample methods to inflict local DNA damage are available.

Our findings support the previously proposed hypothesis that the neurodegenerative phenotype of CS is not caused by defective TCR but by defective RNAPII processing^1,11^. An earlier study proposed that cells lacking CSB or CSA have excessive degradation of ectopically expressed RPB1 due to persistent stalling on unrepaired damages^35^. However, in our local and global DRB assay, we detect the opposite phenotype: RNAPII clearance and degradation are defective in CSB and CSA knockout cells, with stalled RNAPII strongly persisting at the damage sites throughout our experiments. Supporting this notion, several previous studies showed that CSB/CSA-deficient cells fail to degrade RNAPII upon UV treatment, whereas UVSSA-deficient cells do degrade RNAPII^6,10,32^. We also confirm these results here in primary fibroblasts from patients with inherited DNA repair syndromes. By testing our isogenic collection of TCR knockout cell lines side-by-side, we confirmed that only CSB and CSA deficiency causes RNAPII-processing defects, while none of the other TCR factors manifested this phenotype, which is seen in knockout cells and patient-derived fibroblasts. Therefore, this dataset provides key insight into the phenotypical discrepancies observed previously that were associated with deficiencies in different NER genes. The RNAPII-processing defects, particularly observed in CSB and CSA knockout cell lines, correlate well with the neurodegenerative symptoms seen in CS. Correspondingly, deficiencies in downstream TCR factors do not cause defective RNAPII processing, and inactivating mutations in those genes generally manifest as non-neurodegenerative disorders, including UV^S^S or most of XP.

Besides clarification of the discrepancies between clinical phenotypes associated with NER defects, the data presented in this study provide new mechanistic insights into how the processing of damage-stalled RNAPII is regulated. Ubiquitylation of damage-stalled RNAPII involves both proteolytic K48 and non-proteolytic K63 ubiquitin linkages, which are both dependent on CSA^11^ and stimulated by ELOF1 and UVSSA^10^. In our isogenic TCR knockouts, we saw that RNAPII ubiquitylation is completely abolished in CSB^KO^ and CSA^KO^ cells. Interestingly, the residual RNAPII ubiquitylation left in ELOF1^KO^ and UVSSA^KO^ cells was sufficient to induce proteolytic clearance of RNAPII from damaged sites. In agreement with our finding, previous observations showed that UVSSA^KO^ cells have reduced RNAPII ubiquitylation yet degrade RNAPII normally in response to UV damage^6,10^, which we confirm here. These observations suggest that stalled RNAPII is decorated with a variety of ubiquitin linkages resulting in different functional outcomes. Therefore, it will be necessary to develop new tools to detect and manipulate ubiquitin linkages on RNAPII and dissect their roles in defining RNAPII fate^39^.

Rare incidences of CS-like features have been described to be caused by mutations in the downstream NER factors, but these are unlikely to be solely caused by NER defects. The primary reason for this thinking is that defects in XPA, which is essential for downstream TCR, have never been linked to CS-like features except for neurological problems due to potential abnormal repair of base damage. In contrast, the complex combination of symptoms caused by rare mutations in the endonuclease ERCC1-XPF, including, in rare instances, CS-resembling phenotypes, may reflect other functions of this versatile endonuclease outside NER, including interstrand crosslink repair and multiple homologous recombination pathways^30,31^. Moreover, the rare cases of XP/CS and TTD/CS, which are caused by specific mutations in XPG or in the XPB/XPD subunits of TFIIH^27,31^, could either be explained by gain-of-function mutations that lead to impaired RNAPII clearance or by the involvement of these proteins in other process and DNA repair pathways. For example, bi-allelic mutations in the essential DSB repair factor XRCC4 were previously shown to give rise to a CS-like phenotype^40^. Another possibility would be that these rare phenotypes of XP/CS, despite showing normal clearance of RNAPII, are caused by the accumulation of other repair intermediates. Indeed, recent work revealed that the absence of ERCC1-XPF or XPG, but not XPA, leads to persistent binding of TFIIH to non-excised DNA damage, correlating with developmental arrest and neuronal dysfunction in nematodes^41^.

Recent studies proposed an alternative explanation for the discrepancy between CS and other NER phenotypes, suggesting that defects in the transcription-coupled repair of DNA-protein crosslinks (TC-DPC repair) may underly the features of CS^42–44^. This newly discovered pathway repairs bulky DPCs, such as those induced by formaldehyde, in actively transcribed genes in a CSB/CSA-dependent manner. Intriguingly, other TCR factors, including UVSSA, XPA, XPF, and XPG, are not involved in TC-DPC repair, hinting that defective TC-DPC repair is the cause of the neurodegenerative features of CS^42–44^ and an inherited bone marrow failure syndrome, AMeDS, patients of which display CS-like manifestations as well as aplastic anemia, due to the lack of endogenous aldehyde clearance^45^. Effectively, we observe a similar correlation in this study with RNAPII processing, where defects in the processing of damage-stalled RNAPII match CS protein deficiency. The correlation between these two separate observations poses the question of the nature and source of endogenous DNA lesions in CS patients that make CSB or CSA deficiency so detrimental. Although we mainly focused on UV-induced lesions in this study, we speculate that the endogenously occurring DNA lesions, which cause RNAPII stalling, could be a combination of substrates for either TC-DPC repair (e.g., aldehydes) and TCR (e.g., reactive oxygen species-induced cyclopurines). Both types of lesions cause RNAPII stalling and require efficient RNAPII processing executed by CSB and CSA, which subsequently funnel into distinct repair pathways to remove the different types of DNA lesions. Therefore, we hypothesize that it is not the defective repair of either pathway but the defective RNAPII processing in both pathways that causes the neurodegenerative features of CS.

In conclusion, using a new assay to track the fate of elongating RNAPII at sites of UV-induced DNA lesions, we discovered that only CS cells, not WT or non-CS cells, are selectively defective in RNAPII clearance. Hence, our findings provide evidence that defective RNAPII processing and prolonged transcription arrest after DNA damage, rather than compromised DNA repair, may underlie the CS neurodegenerative phenotype.

---

**Fig S1.**
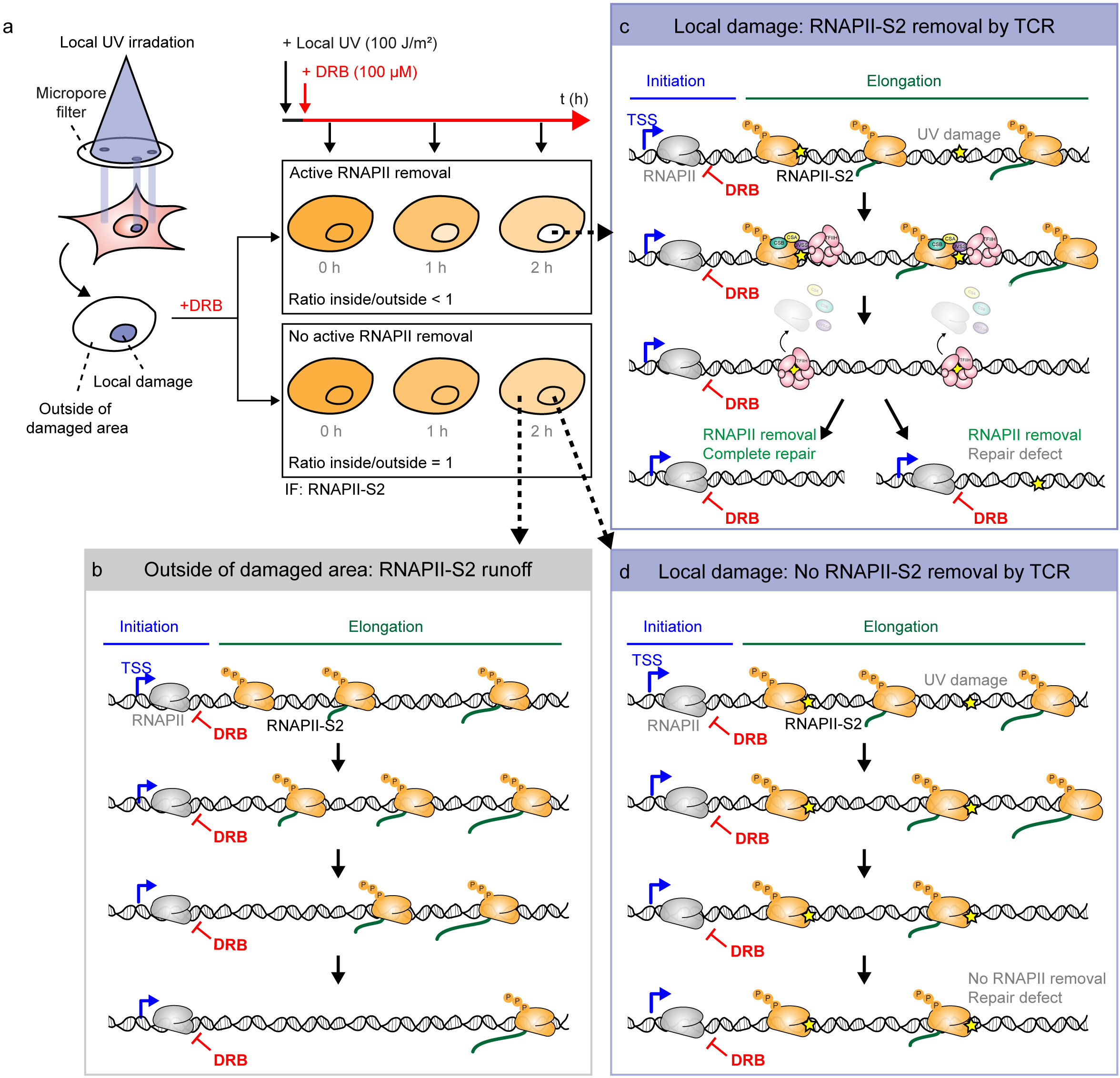
A local UV DRB run-off assay to detect elongating RNAPII clearance. **a.** Experimental set-up with local UV-C irradiation (100 J/m^2^) through 5 µm pores, immediately followed by DRB treatment (100 µM). Cells are fixed at 0, 1, or 2 h after DRB treatment and stained for CPD and RNAPII-S2. **b.** The DRB treatment inhibits new RNAPII molecules from entering gene bodies, causing an RNAPII run-off and an overall decrease in RNAPII-S2 signal. The overall decrease is measured outside sites of local damage. **c.** Inside regions of local UV damage, RNAPII can run-off or stall at a DNA lesions and be removed. The signal inside local damage is the sum of run-off and clearance. **d.** If RNAPII is not removed from DNA lesions, then the signal inside regions of local damage will remain similar to those outside local damage.

**Fig S2.**
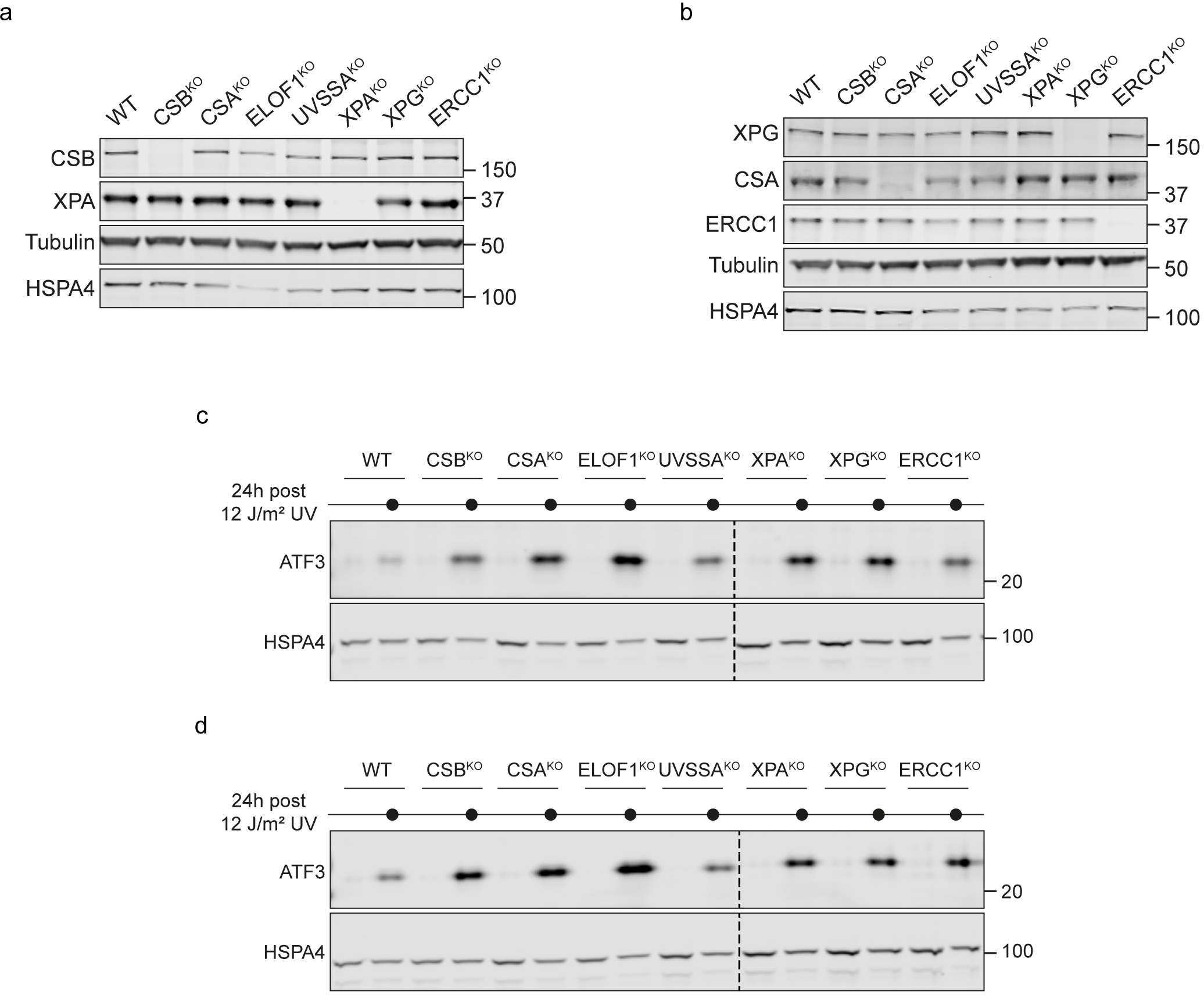
Validation of isogenic collection of TCR knockout cells and their defect in transcription recovery. **a-b.** Validation of the TCR knockouts by western blotting. The indicated antibodies were used to analyze whole cell lysates from parental and knockout RPE1 cell lines. **c-d.** Biological replicates of ATF3 protein levels detected by western blot from whole cell lysates of RPE1 WT and the indicated isogenic TCR knockouts. Cells were either kept unirradiated or exposed to 12 J/m^2^ UV-C and collected after 24 h. The HSPA4 signal was used as a loading control.

**Fig S3.**
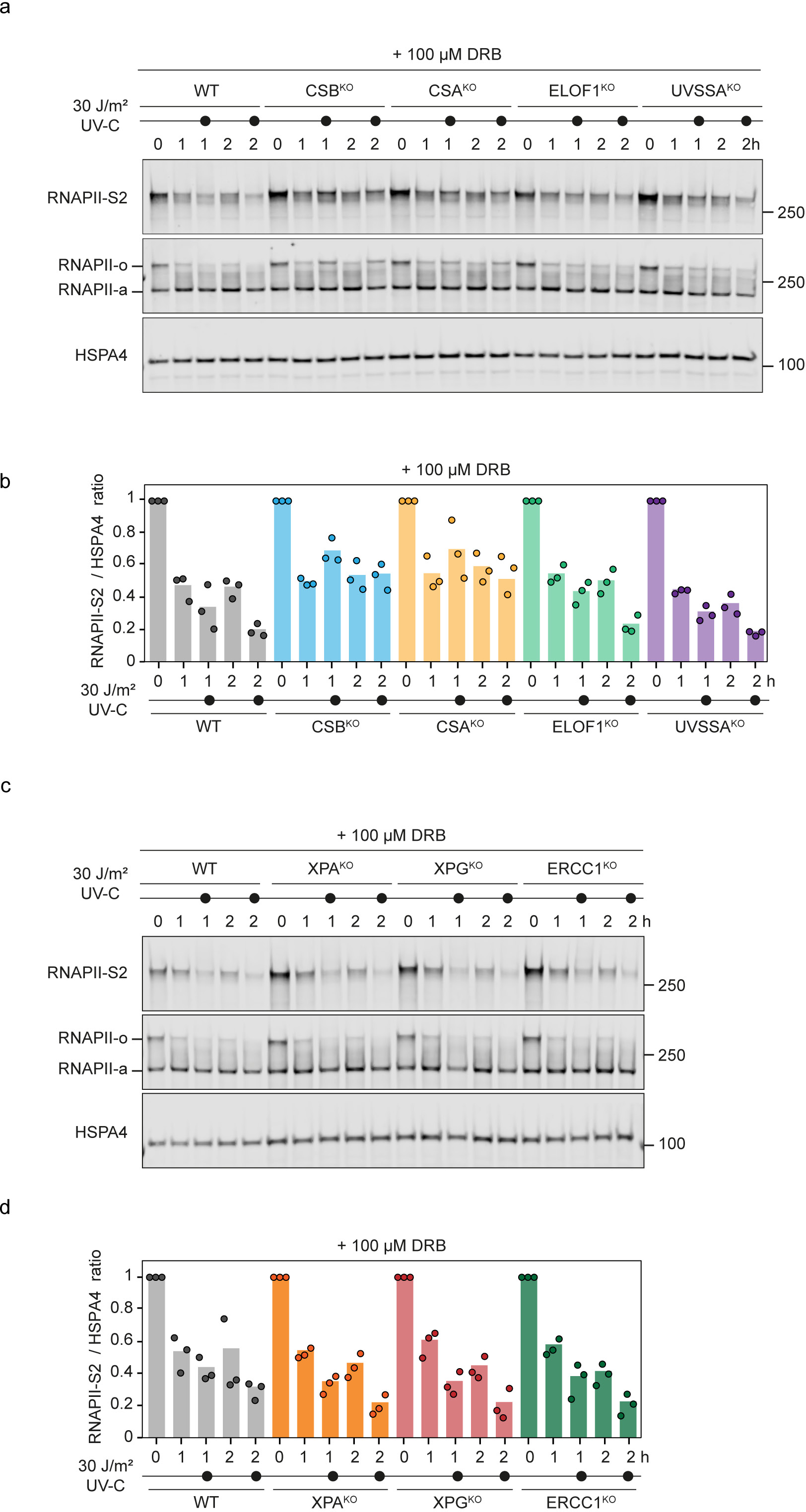
Global degradation of damage-stalled RNAPII in TCR knockout cells. **a, c.** Representative images of the RNAPII signal detected by western blot analysis from whole cell lysates of RPE1 WT and the indicated TCR knockout cells at 0, 1, and 2 h after global UV damage (30 J/m^2^). Cells were treated with 100 μM DRB immediately after UV irradiation to prevent new RNAPII molecules from moving into gene bodies. An antibody against phosphorylated RNAPII detects only the elongating form RNAPII-S2 (upper blot). An antibody targeted against the N-terminal domain of RNAPII detects both initiating (RNAPIIa) and the elongating (RNAPIIo) forms (middle blot). Note that RNAPII-S2 and RNAPIIo represent the same pool of phosphorylated RNAPII. Different abbreviations were used to distinguish between the signals detected by the different antibodies. The HSPA4 signal was used as a loading control (lower blot). **b, d.** Quantification of the blots in **a, c**. The RNAPII-S2 mean pixel intensity was divided by the mean pixel intensity of HSPA4. Then, the value of each time point and condition was normalized to 0 h. A ratio below 1 in unirradiated cells indicates a general RNAPII-S2 run-off caused by DRB. The additional drop in the ratio in UV-treated cells indicates an active RNAPII-S2 degradation by TCR. The bar represents the mean of three independent biological replicates. The individual means of the three biological replicates are depicted as solid circles with black strokes.

**Fig S4.**
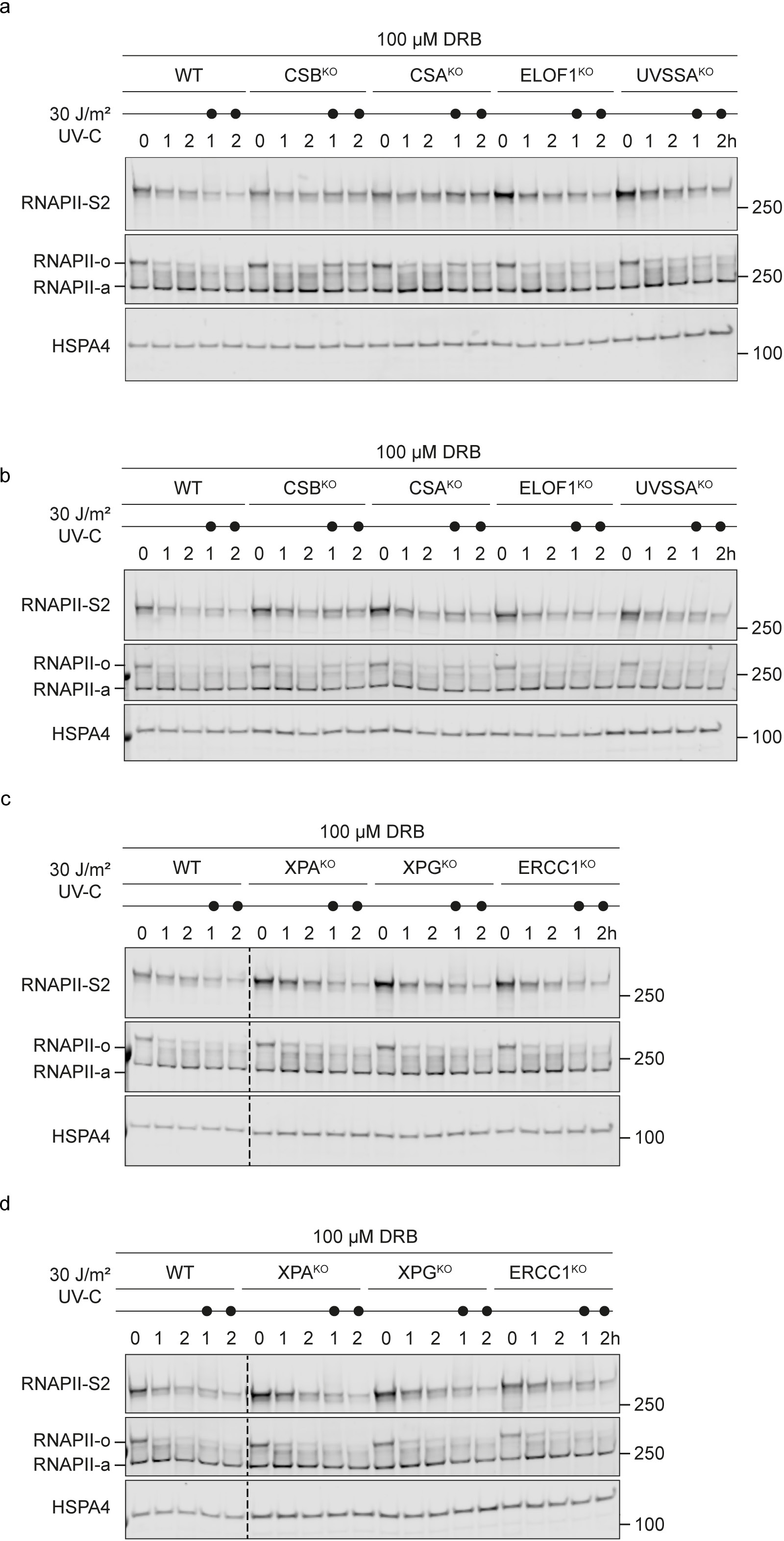
Replicates of the global DRB run-off assay in TCR knockout cells. **a-d.** Biological replicates of the RNAPII signal detected by western blot analysis from whole cell lysates of RPE1 WT and the indicated TCR knockout cells at 0, 1, and 2 h after global UV damage (30 J/m^2^). Cells were treated with 100 μM DRB immediately after UV irradiation to prevent new RNAPII molecules from moving into gene bodies. An antibody against phosphorylated RNAPII detects only the elongating form RNAPII-S2 (upper blot). An antibody targeted against the N-terminal domain of RNAPII detects both initiating (RNAPIIa) and the elongating (RNAPIIo) forms (middle blot). Note that RNAPII-S2 and RNAPIIo represent the same pool of phosphorylated RNAPII. Different abbreviations were used to distinguish between the signals detected by the different antibodies. The HSPA4 signal was used as a loading control (lower blot).

**Fig S5.**
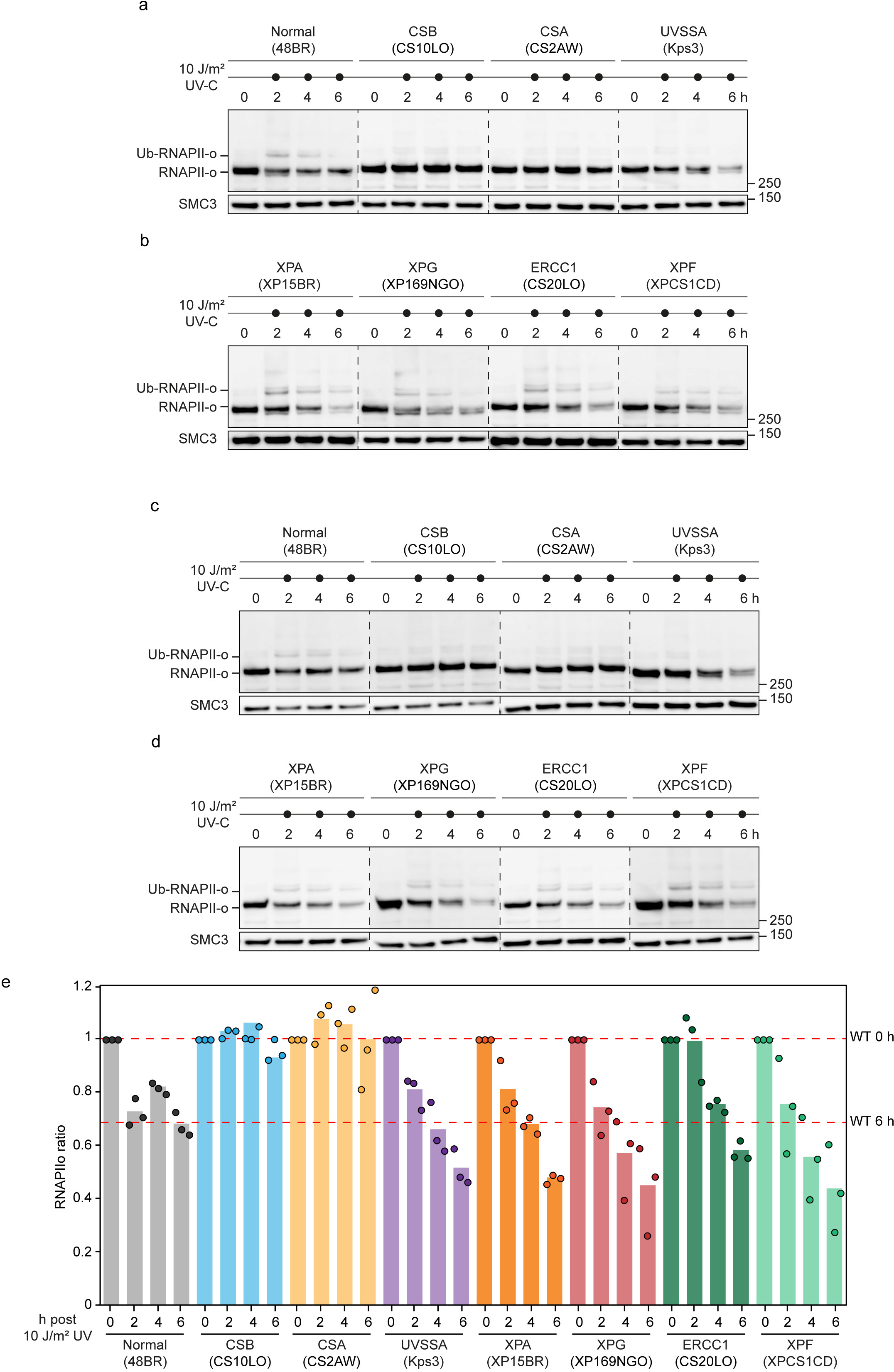
Replicates of ubiquitylation and global degradation of damage-stalled RNAPII in patient-derived cells. **a-d.** Biological replicates of the RNAPII signal detected by western blot analysis from whole cell lysates of patient-derives normal and the indicated TCR mutant cells at 0, 2, 4, and 6 h after global UV damage (10 J/m^2^). Cells were treated with 100 μM cycloheximide 1h before UV irradiation and incubated in the same media after UV exposure. An antibody against hyperphosphorylated C-terminal domain (CTD) Ser2 (H5) was used to detect the elongating RNAPIIo. The SMC3 signal was used as a loading control. **e.** Quantification of the blots in **a-d** and figure 6 a-b. The total RNAPIIo mean pixel intensity was divided by the mean pixel intensity of SMC3. Then, the value of each UV-treated time point was normalized to its respective 0 h condition. Here, a ratio below 1 indicates an active RNAPII-S2 degradation by TCR. The bar represents the mean of three independent biological replicates. The individual means of the three biological replicates are depicted as solid circles with black strokes.

## Acknowledgments

We thank Annelotte P. Wondergem for generating XPA^KO^ cells. MSL laboratory was supported by the Netherlands Scientific Organization (ENW grant OCENW.M20.056, and VIDI grant ALW.016.161.320) and the European Research Council Consolidator Grant STOP-FIX-GO (grant agreement No 101043815). TO laboratory was supported by Practical Research Project for Rare / Intractable Disease (JP24ek0109765 to YN; JP23ek0109678 to TO), grants in Aid for Scientific Research KAKENHI from the Japan Society for the Promotion of Science (JP24K02223 to YN; JP23H00516 to TO), the JST FOREST Program (JPMJFR221E to YN). The funders had no role in study design, data collection and analysis, the decision to publish, or the preparation of the manuscript.

## Data Availability

This published article (and its supplementary information files) includes all data generated or analyzed during this study.

## Author contribution

PJvdM performed all the local DRB run-off assays, incision assays, and detection of Ub-RNAPIIo in RPE1 cells. GY performed all the global DRB run-off assays, ATF3 activation, and validation of the isogenic collection of TCR knockout cell lines. YN performed all the Ub-RNAPIIo detection in patient-derived cell lines. PJvdM, GY, YN, TO, and MSL interpreted the data and wrote the manuscript.

## Methods

### Cell lines and cultures

The RPE1 cell lines were cultured at 37°C in an atmosphere of 5% CO2 in DMEM GlutaMAX (Thermo Fisher Scientific) supplemented with penicillin/streptomycin (Sigma), and 8% fetal bovine serum (FBS; Bodinco BV or Thermo Fischer Scientific (Gibco)). The patient-derived cell lines were maintained in DMEM (WAKO) supplemented with 10% FCS (Hyclone) and antibiotics unless otherwise noted. All cell lines are listed in Supplementary Table 1.

### Generation of knockout cells

Parental RPE1-hTERT cells stably expressing inducible Cas9 (iCas9) that are also knockout for *TP53* and the puromycin-N-acetyltransferase *PAC1* gene were described previously (referred to as RPE1-iCas9)^8^. RPE1-iCas9 cells were transfected with Cas9-2A-EGFP (pX458; Addgene #48138) containing a gRNA against CSB, CSA, ELOF1, UVSSA, XPA, XPG, or ERCC1 from the TKOv3 library using lipofectamine 2000 (Invitrogen). The sgRNAs are listed in Supplementary Table 2, and plasmids in Supplementary Table 3. Cells were FACS sorted on EGFP and plated at low density, after which individual clones were isolated, expanded, and verified by western blot analysis and/or Sanger sequencing using the oligos listed in Supplementary Table 4.

### Western blotting

Total cell lysates were harvested by scraping cells in the Laemmli-SDS sample buffer. Chromatin fractions were obtained by lysing cell pellets on ice in EBC-1 buffer (50 mM Tris [pH 7.5], 150 mM NaCl, 2 mM MgCl2, 0.5% NP-40, and protease inhibitor cocktail (Roche)) for 20 min at 4° C, on a rotating wheel followed by centrifugation and removal of the supernatant. Chromatin pellets were resuspended in the Laemmli-SDS sample buffer. Total cell lysates or chromatin fraction samples were boiled for 10 min at 95°C. Proteins were separated on Criterion™ XT Tris-Acetate 3–8% Protein Gels (BioRad, #3450131) in Tris/Tricine/SDS Running Buffer (BioRad, #1610744) or on Criterion Xt bis-tris 4-12% gels in MOPS running buffer. Then, blotted onto PVDF membranes (IPFL00010, EMD Millipore) in Tris/glycine blotting buffer (0.025 M Tris, 0.192 M glycine) with 20% methanol. Membranes were blocked with 5% fat-free milk in PBS with 0.1 % Tween-20 for 1 h at room temperature. Membranes were then probed with indicated antibodies in 5% fat-free milk in PBS with 0.1 % Tween-20 (Antibodies are listed in Supplementary Table 5). Proteins were stained with fluorochrome-conjugated secondary antibodies and were detected on an Odyssey CLx system and Image Studio software (Li-Cor).

### Incision assay (γH2AX after trabectedin)

Cells were treated with 10 nM trabectedin (MedChemExpress) for 4 h. During the last 15 min, 20 µM 5-Ethynyl-2’-deoxyuridine (5-EdU; Jena Bioscience) was added. Cells were then fixed with 3.7% formaldehyde in PBS for 15 min at room temperature. Cells were subsequently permeabilized with 0.5% Triton X-100 in PBS for 10 min at room temperature and blocked in 3% bovine serum albumin (BSA, Thermo Fisher) in PBS. Dividing cells were visualized by Click-IT chemistry, labeling the cells for 30 minutes with a mix of 6 µM Atto azide-Alexa594 or Atto azide-Alexa647 (Atto Tec), 4 mM copper sulfate (Sigma) and 10 mM ascorbic acid (Sigma) in a 50 mM Tris-buffer (pH 8). After washing with PBS, cells were blocked with 100 mM glycine (Sigma) in PBS for 10 min at room temperature and subsequently with 0.5% BSA and 0,05% tween 20 in PBS for 10 minutes at room temperature. To visualize γH2AX, cells were incubated with a primary antibody for phospho-Histone H2A.X Ser139 (JBW301, Merck) for 2 h at room temperature followed by a secondary antibody Anti-Mouse Alexa 555(A-21424, Thermo Fisher) or Anti-Mouse Alexa 647 (A-21235) and DAPI for 1 h at room temperature and mounted in Polymount (Brunschwig).

### Local DRB-run-off immunofluorescence microscopy assay

Cells were plated in DMEM supplemented with 8% FBS, followed by serum starvation in DMEM without FBS for at least 24 h to reduce the number of replicating cells. Cells were locally UV irradiated through 5 μm pore filters (Milipore; TMTP04700) with 100 J/m^2^ and immediately treated with 100 µM DRB (Sigma, D1916) for the indicated time periods, washed with PBS and fixed with 3.7% formaldehyde in PBS for 15 min at room temperature. Cells were then permeabilized with 0.5% Triton X-100 in PBS for 10 min at room temperature. After washing with PBS, cells were blocked with 100 mM glycine (Sigma) in PBS for 10 min at room temperature and subsequently with 0.5% BSA and 0,05% tween 20 in PBS for 10 min at room temperature. To visualize elongating RNAPII, cells were incubated with a primary antibody for RNAPII-S2 (ab5095, Abcam) and for CPD damages (CAC-NM-DND-001, Cosmo Bio) for 2 h at room temperature and then with secondary antibodies Anti-Mouse Alexa 488 (A-11029, Thermo Fisher) and Anti-Rabbit Alexa 555 (A-21429 Thermo Fisher) and DAPI for 1 h at room temperature. Cells were subsequently mounted in Polymount (Brunschwig).

### Microscopic analysis of fixed cells

Images of fixed samples were acquired on a Zeiss AxioImager M2 widefield fluorescence microscope equipped with 63x PLAN APO (1.4 NA) oil-immersion objectives (Zeiss) and an HXP 120 metal-halide lamp used for excitation. Fluorescent probes were detected using the following filters for DAPI (excitation filter: 350/50 nm, dichroic mirror: 400 nm, emission filter: 460/50 nm), Alexa 555/594 (excitation filter: 545/25 nm, dichroic mirror: 565 nm, emission filter: 605/70 nm), or Alexa 647 (excitation filter: 640/30 nm, dichroic mirror: 660 nm, emission filter: 690/50 nm). Images were recorded using ZEN 2012 software (blue edition, version 1.1.0.0) and analyzed in Image J (1.48v).

### Global DRB run-off assay by western blotting

Cells were cultured in DMEM supplemented with 8% FBS. Then, cells were globally UV irradiated with 30 J/m^2^ and immediately treated with 100 µM DRB (Sigma, D1916) for the indicated time periods. Cells were washed with PBS and total cell lysates were harvested by scraping cells in the Laemmli-SDS sample buffer. Proteins were separated and analyzed by western blotting.

### Detection of the elongating RNAPIIo after UV irradiation in patient-derived cell lines

Cells were cultured in a medium supplemented with 100 μM cycloheximide for 1 h before UV irradiation. Cells were irradiated (10 J/m^2^ of UV-C) and incubated for the indicated time periods in a medium containing cycloheximide. Whole cell lysates were resolved on 6% SDS-PAGE gels and transferred to the PVDF membrane. RNAPIIo was detected with an H5 antibody (recognizing phosphorylated CTD-Ser2).

---

**Supplementary Table 1:** Cell lines.

| Cell lines | Source |
| --- | --- |
| RPE1-iCas9 (WT) | 8 |
| RPE1-iCas9 CSA-KO (3-1) | 8 |
| RPE1-iCas9 CSB-KO (1-15) | 8 |
| RPE1-iCas9 ELOF1-KO (2-16) | 8 |
| RPE1-iCas9 ERCC1-KO (16) | 46 |
| RPE1-iCas9 UVSSA-KO (3-9) | 8 |
| RPE1-iCas9 XPA-KO (1) | This study |
| RPE1-iCas9 XPG-KO (21) | 47 |
| 48BR (WT primary fibroblast) | 48 |
| CS10LO (CS-B primary fibroblast) | 49 |
| CS2AW (CS-A primary fibroblast) | 50 |
| Kps3 (UVSS-A primary fibroblast) | 6 |
| XP15BR (XP-A primary fibroblast) | 51 |
| XP169NGO (XP-G primary fibroblast) | This study |
| CS20LO (ERCC1 primary fibroblast) | 30 |
| XPCS1CD (XP-F primary fibroblast) | 30 |

**Supplementary Table 2:**
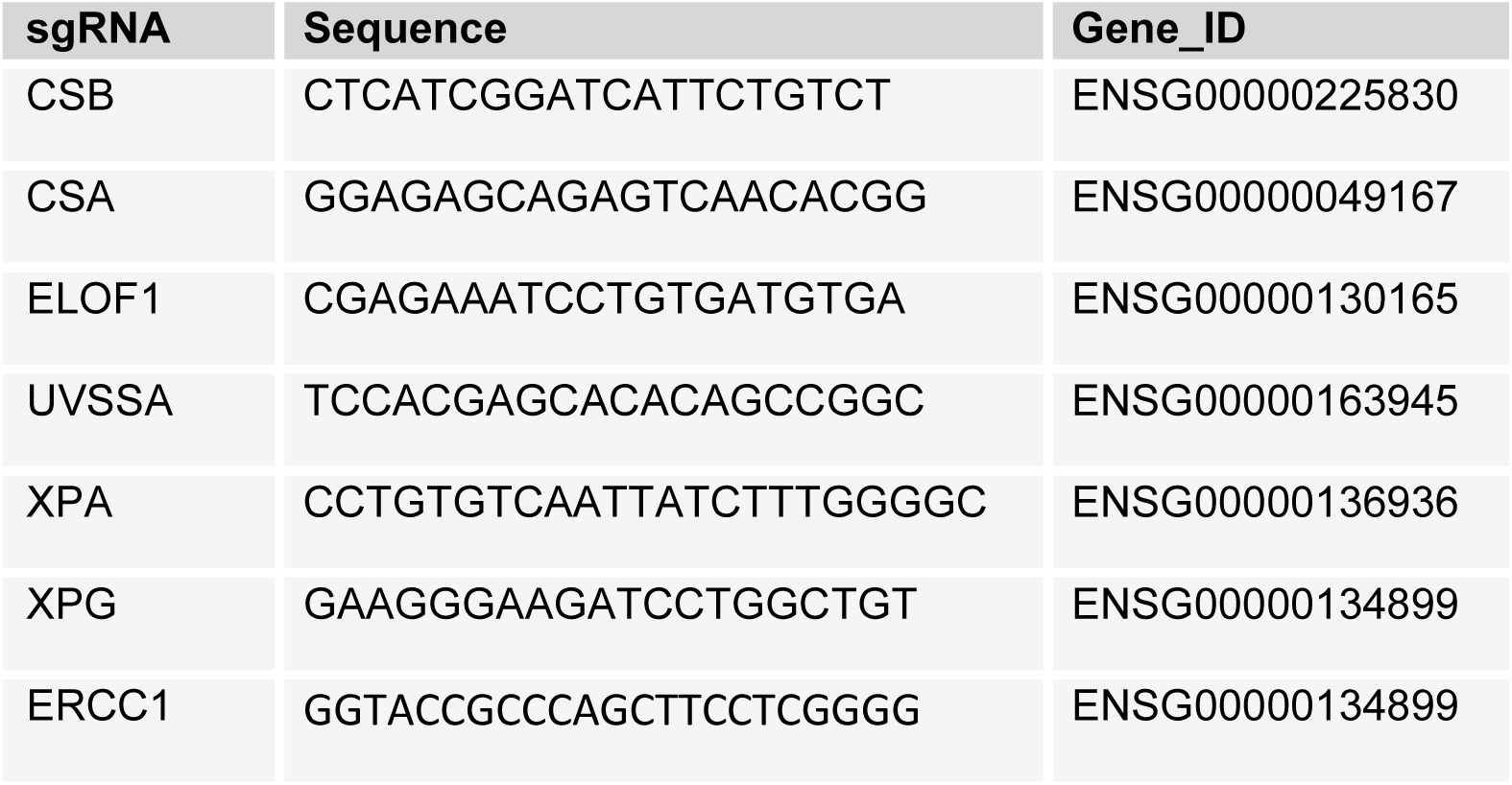
sgRNAs.

**Supplementary Table 3:** Plasmids.

| Plasmid | Origin | Identifier |
| --- | --- | --- |
| pX458 | Addgene #48138 | pML#006 |
| pX458-(Cas9-2A-GFP)-sgCSB-1 | 10 | pML#003 |
| pX458-(Cas9-2A-GFP)-sgCSA-3 | 10 | pML#219 |
| pX458-(Cas9-2A-GFP)-sgELOF1-2 | 10 | pML#186 |
| pX458-(Cas9-2A-GFP)-sgUVSSA_3 | 10 | pML#220 |
| pLV-U6g-PPB-sgXPA_2 | This study | sgML#002 |
| pX458-(Cas9-2A-GFP)-sgXPG | 47 | pML#223 |
| pLV-U6g-PPB-sgERCC1_1 | 46 | sgML#049 |

**Supplementary Table 4:**
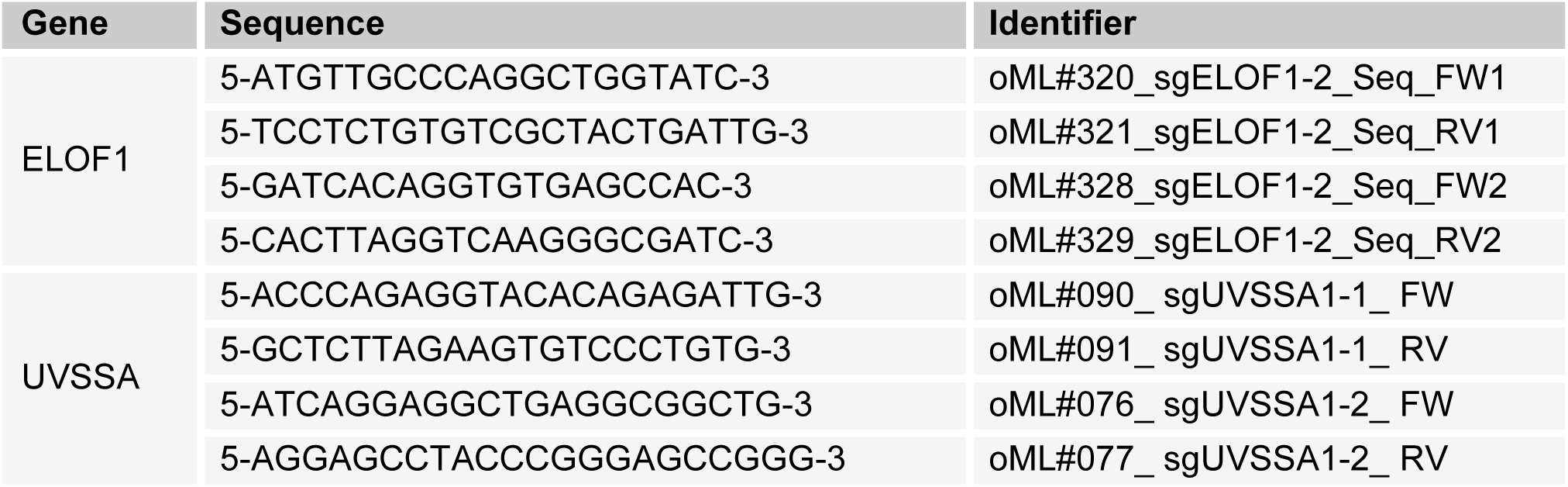
Oligos.

**Supplementary Table 5:**
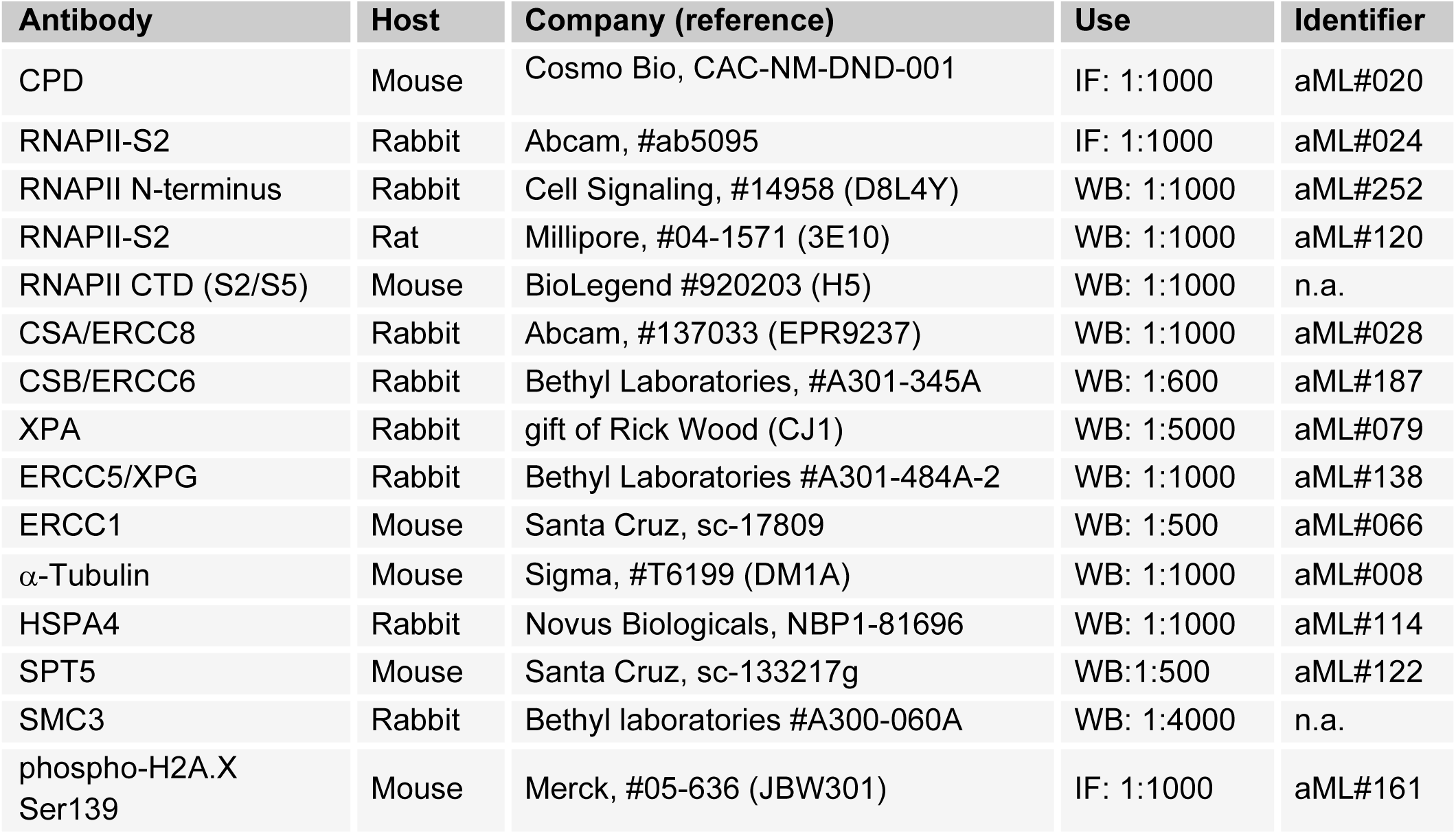
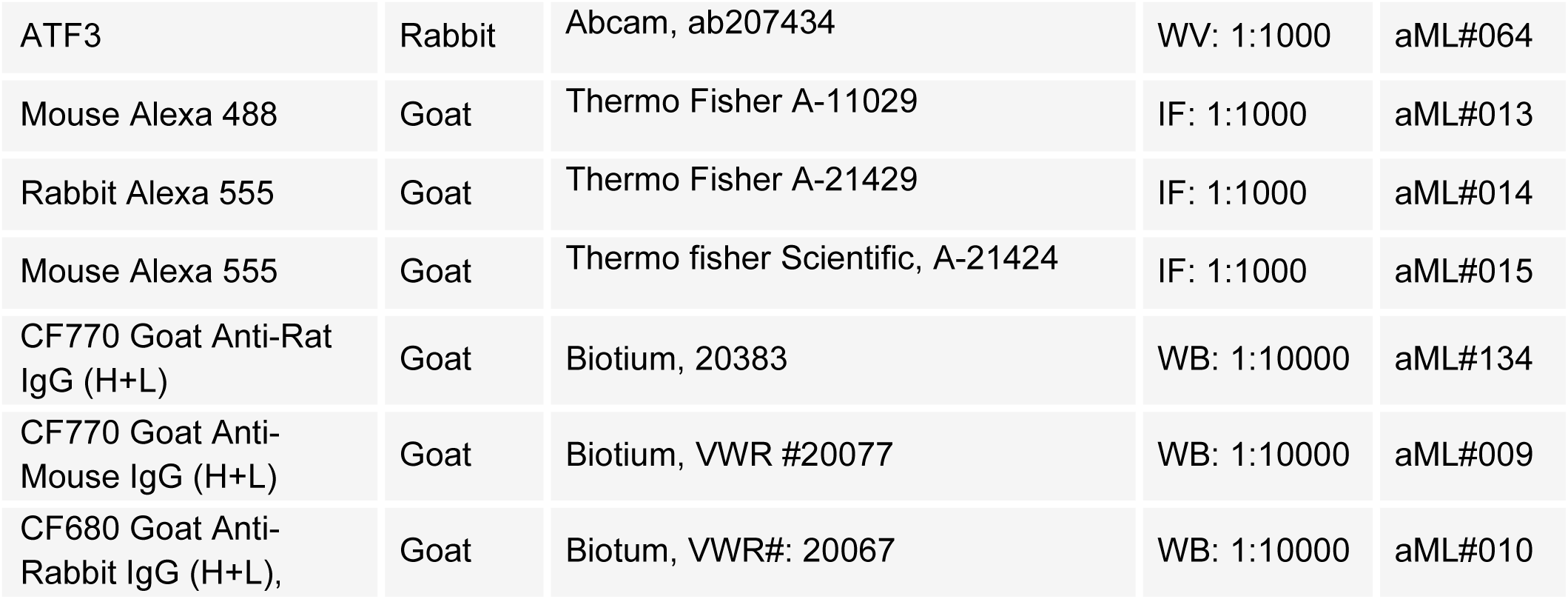
Antibodies.

